# An Artificial Intelligence Workflow for Defining Host-Pathogen Interactions

**DOI:** 10.1101/408450

**Authors:** Daniel H Fisch, Artur Yakimovich, Barbara Clough, Joseph Wright, Monique Bunyan, Michael Howell, Jason Mercer, Eva-Maria Frickel

## Abstract

For image-based infection biology, accurate unbiased quantification of host-pathogen interactions is essential, yet often performed manually or using limited enumeration employing simple image analysis algorithms based on image segmentation. Host protein recruitment to pathogens is often refractory to accurate automated assessment due to its heterogeneous nature. An intuitive intelligent image analysis program to assess host protein recruitment within general cellular pathogen defense is lacking. We present HRMAn (Host Response to Microbe Analysis), an open-source image analysis platform based on machine learning algorithms and deep learning. We show that HRMAn has the capability to learn phenotypes from the data, without relying on researcher-based assumptions. Using *Toxoplasma gondii* and *Salmonella typhimurium* we demonstrate HRMAn’s capacity to recognize, classify and quantify pathogen killing, replication and cellular defense responses.

## Introduction

High content imaging (HCI) has revolutionized the field of host-pathogen interaction by allowing researchers to perform image-based large-scale compound and host genome-wide depletion screens in a high-throughput fashion (1, 2). The majority of these screens assess host-pathogen interactions using bulk colorimetric or automated enumeration of pathogen growth at the population level (3). Additionally, quantification of host-pathogen interaction (e.g. analysis of host protein recruitment to the pathogen) in general is often performed manually. However, to meaningfully dissect cell-mediated pathogen control, it is imperative to quantify the host response and pathogen fate at the single-cell level. Many open-source (e.g. CellProfiler (4)) and proprietary (e.g. Perkin Elmer Harmony) analysis software packages have been developed and successfully employed for exactly this purpose (5–7). To advance the state of the art in image analysis of host-pathogen interaction, incorporation of cutting-edge machine intelligence algorithms (8, 9) to stratify the image content without the requirement to program complex integrations is needed. HRMAn relies on the same well-established image segmentation strategies as many other programs, but offers an intuitive integration of deep learning for more complex image analysis (an overview of currently available programs can be found in Table S1). Deep neural network-based machine intelligence methods have brought about a revolutionary advance in the field of computer vision, by allowing for machine learning of complex morphologies in a highly generalized fashion (8, 10). These methods have not yet been adapted for the field of host-pathogen interaction. Typically, HCI based fluorescent imaging data from a host-pathogen interaction experiment is analyzed by classical image segmentation (11–14). Occasionally segmentation combined with machine learning based on calculated features has been employed (15). Most of these analysis pipelines make use of open-source programs tailored with additional coding by the user to suit their specific needs. They are not published in their final form as a universal open-source solution. Most importantly, these classical image segmentation and machine learning analysis methods fail at the level of quantifying host protein recruitment to the pathogen. This is largely due to the fact that traditional algorithms quantify phenotypes in a rule-based manner, using bulk statistical properties of microscopy images or their segments. Conversely, deep neural networks make use of complex patterns (e.g. shapes) within the dataset to learn phenotypes and their diversity. The neural network derives these complex patterns in an unsupervised fashion from expert-labeled data. Thus, using pattern complexity to refine classification (10), deep neural networks improve the biological relevance of the phenotypic readouts. While some proprietary solutions have been employed to extract host protein recruitment data, these solutions are impractical for most researchers as they are tied to single and expensive microscopes (16). To date, for the analysis of host protein recruitment to pathogens, artificial intelligence-driven automated analysis is neither available as an open-source or commercial package. Thus, there remains a need for an open-source, intuitive, multi-parametric, flexible, and trainable host-pathogen interaction analysis software that performs at the level of human analytic capacity (17, 18). Here we present a high-throughput, high-content, single-cell image analysis pipeline that incorporates machine learning and a deep convolutional neural network (CNN) ensemble for Host Response to Microbe Analysis (HRMAn; www.hrman.org). To assure its broad applicability to infection biology, HRMAn is based on the data handling environment Eclipse-KNIME (19). The analysis relies on training of machine learning algorithms and deep neural networks that can be tailored to individual researchers’ needs.

## Results

### Architecture of the high-content image analysis pipeline for analyzing host-pathogen interaction

The HRMAn pipeline (Fig. 1), is designed to work with all file types acquired on any HCI platform or fluorescence microscope. Plate maps including experimental layouts, sample groups and replicates can be loaded, enabling HRMAn to automatically cluster results and perform error analysis. Once fed into the HRMAn pipeline, images are automatically preprocessed and clustered based on user-defined parameters (i.e. imaging specifications) and corrected for illumination. The subsequent image analysis proceeds in two stages: in stage 1, HRMAn segments images into pathogen and cell features for single cell analysis. It then classifies these features using a decision tree learning algorithm previously trained on an annotated dataset. In stage 2, HRMAn analyzes cell features associated with the pathogens using a CNN (based on AlexNet architecture) trained to distinguish complex phenotypic patterns of host-protein recruitment (8). Robust classification is achieved by passing convolutions of segmented areas of interest through multiple non-linear filters to identify characteristic phenotypic details. Finally, data is output as a single spreadsheet file providing the researcher with ≥ 15 quantitative descriptions of a pathogen and its interaction with host factors at population and single cell levels (Fig. 1; Readouts). Importantly, by separating the analysis, HRMAn offers researchers the flexibility to perform fast, simple quantitative analysis of infection parameters using stage 1, without analyzing host protein recruitment.

**Fig. 1.**
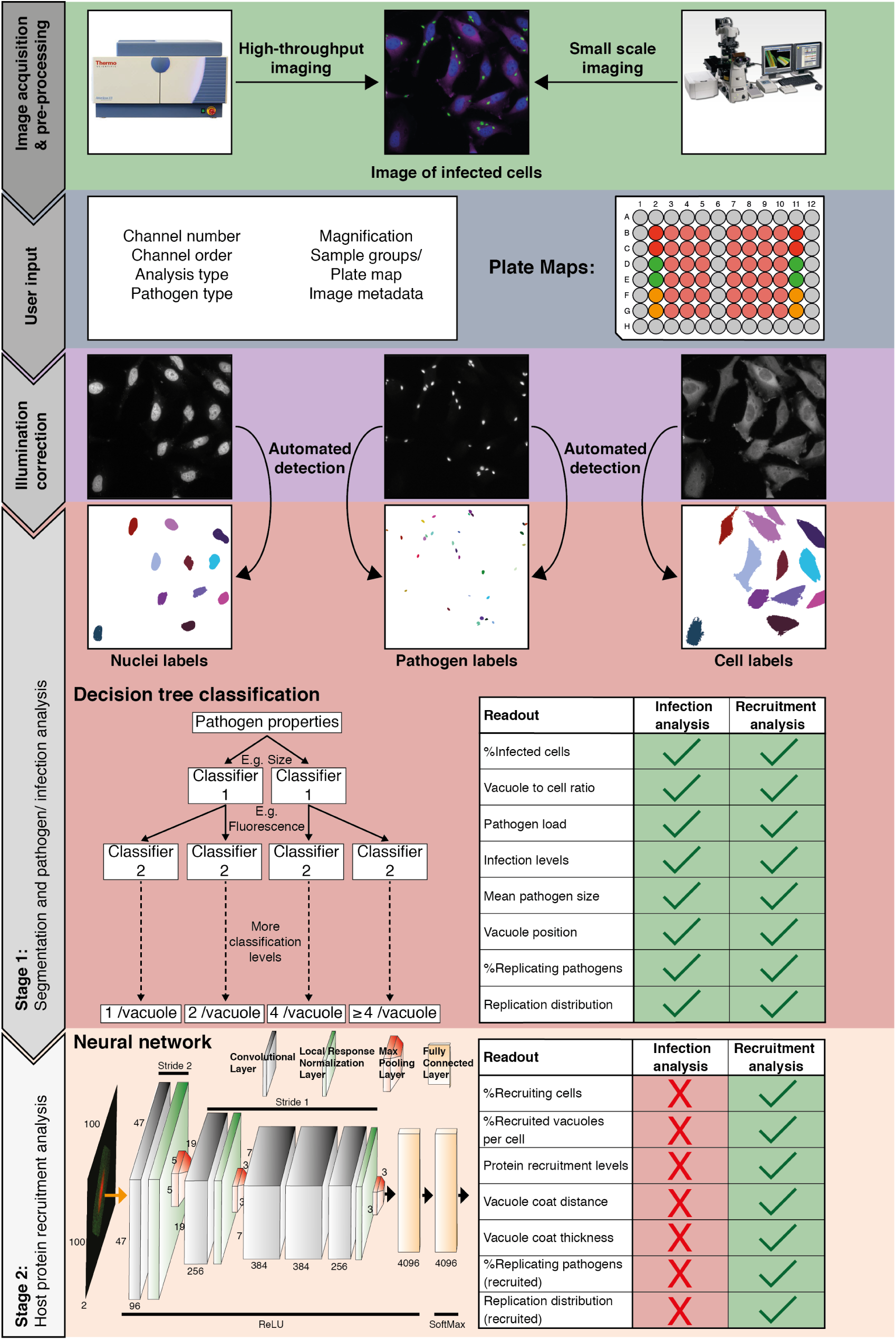
Overview of the HRMAn pipeline. Following image acquisition, on a high-content imaging platform or any other fluorescence microscope, the images can be loaded into the HRMAn software. First, the data is pre-processed and clustered based on user-defined parameters and provided plate maps. Images then undergo illumination correction and automated segmentation. Segmented images are used by a deep convolution neural network (CNN) and other machine learning based algorithms to analyze infection of cells with intracellular pathogens. Depicted is the CNN diagram representing three-dimensional convolutional filters with respective width, height and depth designated on filters facets. Respective change of stride in the groups of hidden layers is depicted above the diagram, while respective activation functions below the diagram. Finally, the data is written as a single file and will provide the researcher with more than 15 different readouts that describe the interaction between pathogen and host cell during infection. HRMAn is based on the open-source data handling environment KNIME making it modular and adaptable to a researcher’s needs. The analysis is based on training of the machine learning algorithms generating high flexibility, which can be tailored to the needs of the user.

### Machine learning and a convolutional neural network drives classification of pathogen replication and host defense

To train for detection and analysis of host-pathogen interactions, HRMAn was provided an annotated dataset of cells infected with an eGFP-expressing version of the parasite *Toxoplasma gondii (Tg)* and stained for various cell features (Fig. 2a) (20, 21). For stage 1 pathogen detection and enumeration training, over 35,000 *Tg*-vacuoles were analyzed by decision tree, gradient boosted tree, and random forest machine learning classification algorithms and cross-validated (Fig. 2b). As each performed equally, a simple decision tree with Minimum Description Length (MDL) pruning, to limit overfitting, was employed for speed and accuracy of pathogen detection (> 99.5% for Tg). Using these parameters, in addition to the readouts from stage 1 (see Fig. 1), HRMAn detected and quantified *Tg*-containing vacuoles harboring 1, 2, 4 or ≥ 4 fluorescent *Tg* (Fig. 2c).

**Fig. 2.**
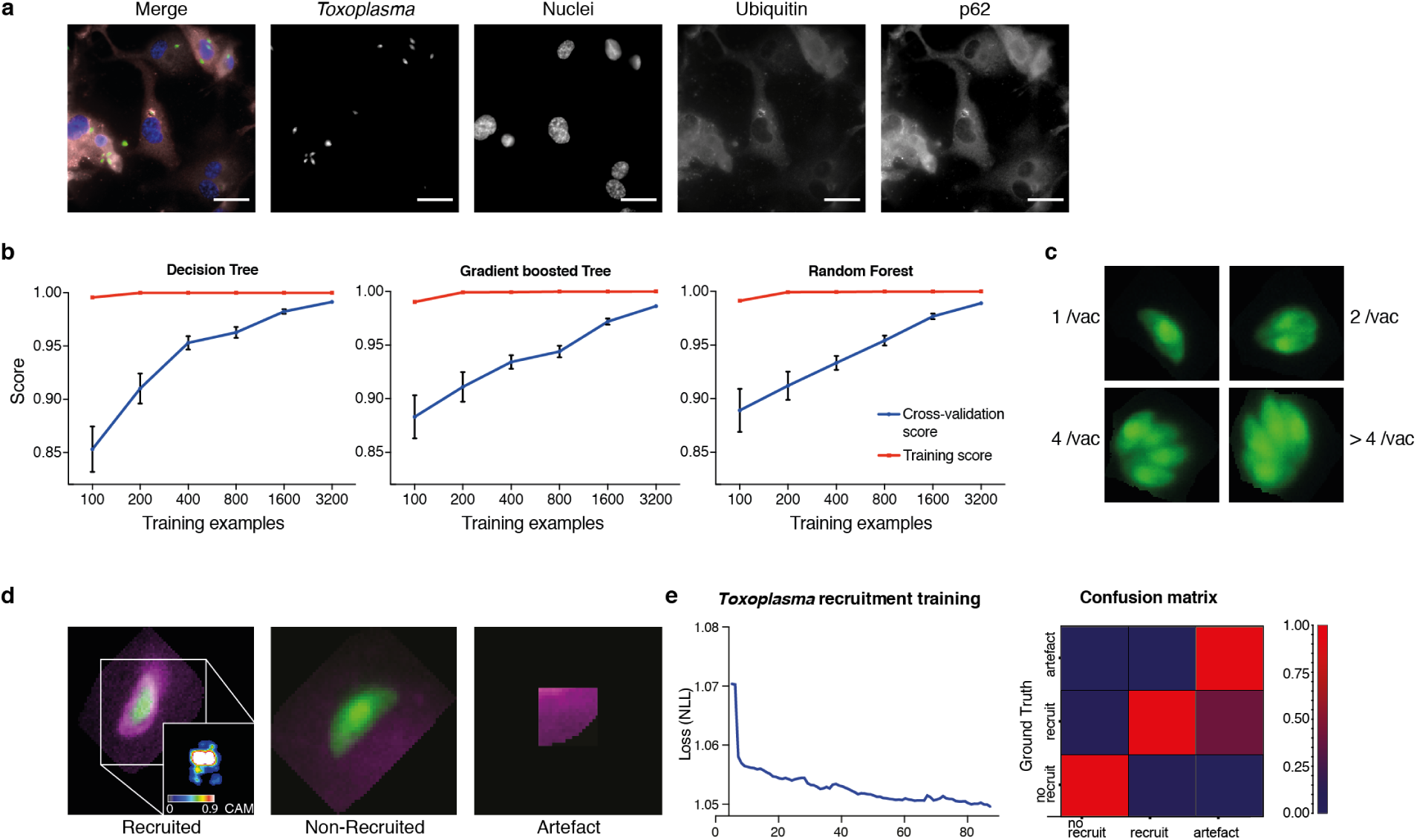
Decision-tree and convolutional neural network training for pathogen replication and host defense protein recruitment analysis. (a) Example images from one field of view. A composite image of all channels (blue: nuclei, green: Tg, red: Ubiquitin, grey: p62) and the single channel images are shown. Scale bar indicates a distance of 30 m. (b) Training and cross-validation of different machine learning classification algorithms to predict parasite replication. (c) Example images of different vacuoles with the resulting classification of a trained decision tree classifier. (d) Resulting classification of the trained deep convolution neural network (CNN) with example vacuoles. For the recruited classification a class activation map (CAM) is depicted to illustrate the focus of the CNN. (e) Decrease of negative log likelihood (NLL) used as loss function during CNN training over training cycles (epochs) for *Toxoplasma gondii* model (left) and confusion matrix of *Toxoplasma gondii* model validation illustrating classification accuracy of labelled data unseen by the model, classification accuracy (0 to 1) during validation is color-coded blue to red (right).

For stage 2, host protein recruitment, the CNN was trained for ubiquitin and p62 recruitment using segmented *Tg* vacuoles defined in Stage 1. Robust classification of host protein recruitment was achieved by passing these regions of interest through multiple non-linear filters to identify and differentiate between no recruitment, recruitment, and analysis artefacts (Fig. 2d). Using backpropagation over 80 training cycles (epochs) with negative log likelihood as a loss function, the deep CNN achieved 92.1% classification accuracy, confirmed by expert based cross-validation (Fig. 2e).

### HRMAn allows for accurate high-throughput analysis of the host defense response to Toxoplasma

To demonstrate the ability of HRMAn and to expand how researchers define and classify host-pathogen interactions, the impact of IFNγ on *Tg* replication and ubiquitin/p62 recruitment to *Tg* vacuoles was analyzed (Fig. 3). To assure that uninvaded *Tg* parasites do not skew the data, stringent synchronization of infection by centrifugation and washing procedures were employed. In a pilot ds experiment (Fig. S1), staining with the *Tg* vacuole marker GRA2 (Fig. S1a+b) revealed that more than 98% of all parasites captured in the images have successfully invaded and established a PV, irrespective of the *Tg* strain used for infection (Fig. S1b). At the multiplicity of infection (MOI) of 3 used for experiments, this resulted in up to 90% type I and 80% type II *Tg* infected cells (Fig S1c). Importantly, we often observed that a single cell can contain more than one PV.

**Fig. 3.**
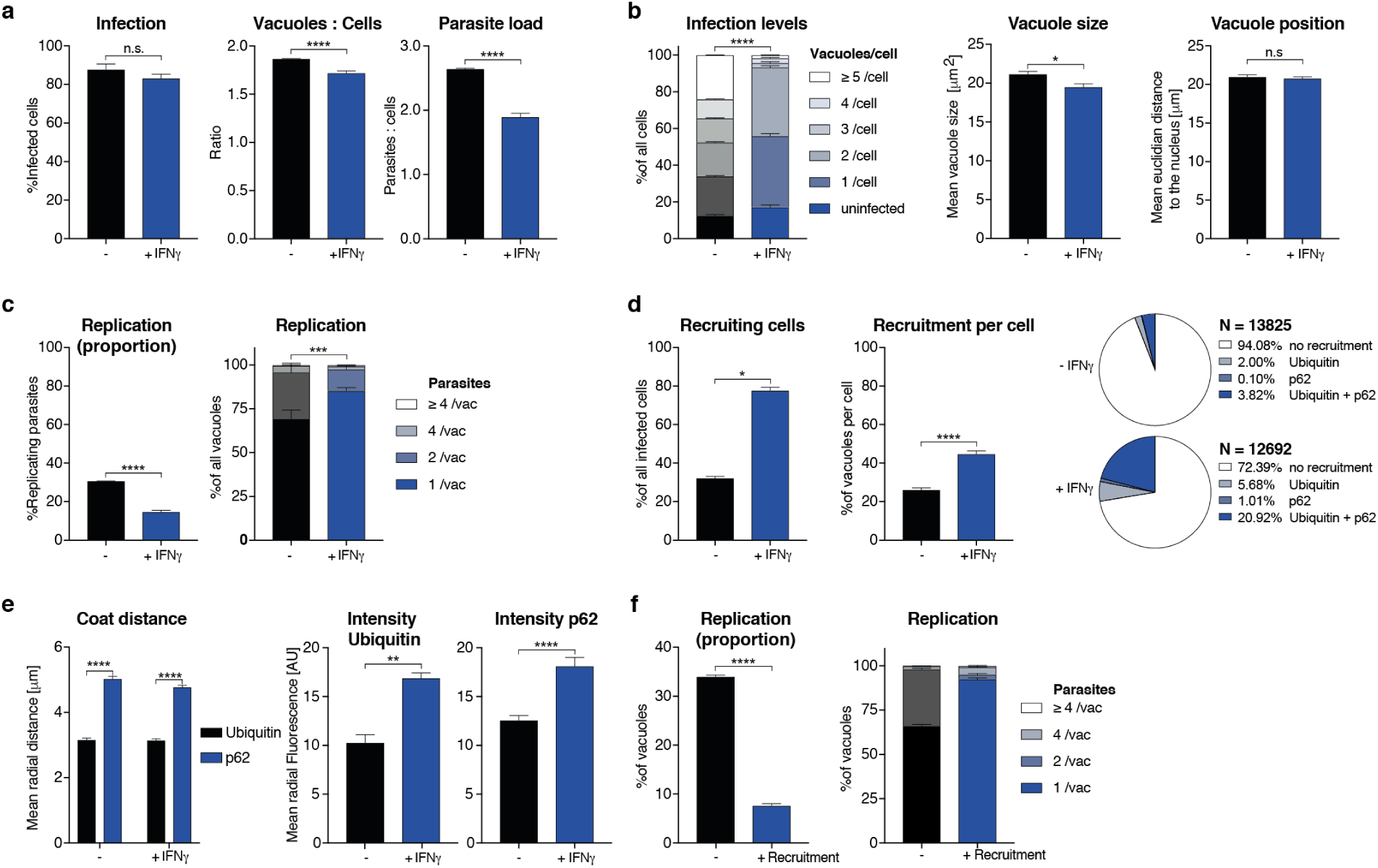
Analysis of *Toxoplasma gondii* infection in IFNγ-treated HeLa cells. HeLa cells were stimulated with 100 IU/mL IFNγ, infected with type I (RH) *Toxoplasma gondii (Tg)* and analyzed 6 hours post-infection. (a) Infection parameters depicted as total percent of *Tg* infected cells, the ratio of *Tg* vacuoles to cells and the ratio of parasites to cells. (b) Cellular readouts showing the proportion of cells that contain a varying numbers of parasite vacuoles, the mean vacuole size of *Tg* and the vacuole position as the value of the mean euclidian distance of *Tg* vacuoles to the host cell nucleus. (c) Replication capacity of *Tg* shown as the proportion of replicating parasites and the distribution of replicating Tg. (d) Cellular response to infection with *Tg* measured as the percentage of cells that decorate vacuoles and the average proportion of vacuoles per cell that are being decorated simultaneously and the overall proportion of ubiquitin and/or p62 decorated *Tg* vacuoles. N shows the total number of vacuoles analyzed for each condition, percentages are indicated in the legend. (e) Properties of the host protein coat on *Tg* vacuoles as the average coat distance for ubiquitin and p62 to *Tg* and mean fluorescence intensity of ubiquitin and p62 at *Tg* vacuoles. (f) Fate of *Tg* vacuoles grouped based on host protein recruitment. The proportion of replicating parasites and the replication distribution based on recruitment status of the vacuole are shown. All data shown above represents the mean of N = 3 experiments *±* SEM. Significance was determined using unpaired t-tests, n.s. = not significant, * p *≤* 0.0332, ** p *≤* 0.0021, *** p *≤* 0.0002 and **** p *<* 0.0001.

Previous reports indicate that HeLa cells restrict the growth of *Tg* through ubiquitination of parasitophorous vacuoles and subsequent non-canonical, p62-dependent autophagy (20, 22). HeLa cells infected with EGFP *Tg* ± IFNγ were fixed 6 hours post-infection (hpi) and stained with Hoechst (nuclei) and antibodies directed against ubiquitin and p62. Overall, 1,350 4-color images were acquired on an automated microscope and loaded into HRMAn for analysis.

HRMAn automatically detected and analyzed more than 15,000 cells resulting in 15 quantitative outputs of host-pathogen interaction (Fig. 3). Population level readouts from stage 1 indicated that IFNγ treatment did not impact the percentage of infected cells but decreased the number of vacuoles within cells as well as the number of parasites per cell (Fig. 3a). As eGFP fluorescence is lost when parasites are killed, a reduction in the ratio between vacuoles and cells serves as an indirect measurement for parasite killing. At the single cell level, HRMAn found that IFNγ treatment resulted in a significant reduction of vacuoles per cell and a minor reduction in mean vacuole size, without impacting vacuole position (Fig. 3b). Concomitant with this reduction in vacuole size, both the percentage of replicating parasites, and the number of parasites per vacuole were significantly reduced by IFNγ treatment (Fig. 3c). Thus, IFNγ mediated control of *Tg* in HeLa cells involves both parasite killing and restriction of *Tg* replication. Importantly, HRMAn offers a wide range of readouts in stage 1 analysis allowing for more detailed information on the dynamics of infection and clearance than typically seen with manual counting. To allow the user to decide which readouts are best suited to answer their specific research question some redundancy has been purposely built in (e.g. mean vacuole size vs. % Replicating). For example, here we focused mainly on parasites per vacuole and the proportion of infected cells, as opposed to the number of individual vacuoles per cell.

In stage 2, analysis of the >25,000 vacuoles identified in stage 1, showed that the number of cells with ubiquitin/p62-positive vacuoles and the percentage of ubiquitin/p62-positive vacuoles per cell increased with IFNγ (Fig. 3d). Distribution analysis indicated that in untreated cells, only 5.92% of vacuoles were decorated with ubiquitin, p62, or both. This number rose to 27.61% in IFNγ-treated cells, the majority of which (20.92%) were double-positive for ubiquitin/p62 (Fig. 3d). By quantifying the radial fluorescence intensity distribution of these host factors, HRMAn revealed that ubiquitin was more closely associated with *Tg* vacuoles than p62 and that recruitment of both was increased by IFNγ treatment (Fig. 3e). This is in agreement with the notion that p62 binds a ubiquitinated vacuole substrate through its UBA domain (20, 21). Finally, by analyzing vacuoles that recruit ubiquitin/p62, HRMAn indicated that restriction of *Tg* replication occurs in vacuoles decorated with these host defense proteins (Fig. 3f). Collectively, this data indicates that in HeLa cells, IFNγ drives both parasite killing as well as recruitment of ubiquitin/p62 to *Tg* vacuoles, which acts to restrict parasite replication (Fig. 3). The results demonstrate the capacity of HRMAn to provide a quantitative, multi-parametric read-out of host-pathogen interaction at population and single-cell levels.

As a high-throughput, high-content analysis program, HRMAn removes experimental size constraints imposed by manual quantification. To illustrate this, HRMAn was used to systematically analyze the impact of IFNγ treatment on type I and type II Toxoplasma strains in 5 human cell lines: HeLa (cervical carcinoma epithelial), PMA-differentiated THP-1 (macrophage-like), A549 (lung carcinoma epithelia), HFF (primary fibroblasts), and HUVEC (primary endothelial cells) (Fig. S2-S8).

First, stage 1 HRMAn was used to ascertain the impact of varying concentrations of IFNγ (50-500 IU/ml) on *Tg* infection, killing, and replication. (Fig. S2). For each cell line (Fig. S2a), a dose-dependent reduction in *Tg* infection was seen (Fig. S2b). Assessment of the vacuole:cell ratio and mean vacuole size indicated that THP-1s, HFFs, and HUVECs limit infection largely by IFNγ-dependent *Tg* killing, while HeLas and A549s do so by restricting replication (Fig. S2c+d). Quantification of the number of parasites per vacuole indicated that HeLas and A549s acutely restrict type I and type II *Tg* replication at all concentrations of IFNγ(Fig. S3b+c), while THP-1s, HFFs, and HUVECs are far more limited in this capacity (Fig. S3a, d+e).

Next, HRMAn was employed on all 5 cell lines infected with either type I and type II *Tg* ± 100 IU/ml IFNγ and immunostained for ubiquitin and p62. Supplementary figures S4-S8 display the 15 quantitative readouts compiled by HRMAn of 9,000 fields of view (90 GB) and >175,000 vacuoles identified in stage 1. Taking advantage of the large-scale capabilities of HRMAn, we found that all cell types can mediate IFNγ-dependent type I and II *Tg* killing (Fig. S4b+c), and growth restriction (Fig. S5a+b) to similar levels. *Tg* vacuoles show strain-dependent (A549, HUVEC), and strain-independent (HFFs) IFNγ-stimulated movement towards the nucleus (Fig. S5c). HRMAn revealed that type II *Tg* grew slower than type I *Tg* in each cell line and that their growth decreased more upon treatment with IFNγ (Fig. S6a-b). Consistent with this, stage 2 HRMAn showed that all cell types could recruit ubiquitin and/or p62 equally well (Fig. S7a), while a greater percentage of type II vacuoles per cell were decorated in response to IFNγ-priming (Fig. S7b). The exception to this were THP-1 cells, which did not mount a strain-specific response (Fig. S7b). Distribution analysis further indicated that THP-1s rather display a higher intrinsic capacity to decorate *Tg* vacuoles than other cell lines, even in the absence of IFNγ (Fig. S7c). While no cell-type dependent differences in ubiquitin or p62 coat distance were observed (Fig. S8a), THP-1s not only decorate vacuoles with more ubiquitin upon IFNγ stimulation, they also appear to recruit p62 in an IFNγ-independent fashion (Fig. S8b). Decorated vacuoles in all cell types displayed a greater ability to restrict the growth of type II versus type I *Tg* upon IFNγ treatment (Fig. S8c-d). These results highlight the ability of HRMAn to provide high-throughput and quantitative single-cell analysis of host-pathogen interactions at a scale not achievable by automated bulk or manual quantification.

### HRMAn can be adapted for bacteria-host interaction analysis

To demonstrate its flexibility, HRMAn was trained to recognize the bacterium *Salmonella typhimurium* (*STm*) - a pathogen 16x smaller than *Tg* (0.5 m vs. 8 m) - and then set to analyze the impact of IFNγ on bacterial killing, replication, and ubiquitin recruitment. Stage 1 outputs showed that similar to *Tg* (Fig. 3), IFNγ treatment in HeLa cells reduced the ratio of *STm* vacuoles/cell and the bacterial load, without impacting the percent of infected cells (Fig. 4a). At the single cell level, HRMAn found a significant reduction in the number of *STm* vacuoles/cell, consistent with a reduction in vacuole size, percent of replicating bacteria, and reduced numbers of *STm*/vacuole (Fig. 4b+c). These results demonstrate that HeLa cells can control infection with *STm* through IFNγ dependent bacterial killing and growth restriction. For stage 2, we used the *Tg* recruitment model as input to retrain HRMAn for quantification of ubiquitin recruitment to *STm* (Fig. 4d). This allowed us to achieve 69.9% classifica-accuracy, confirmed by expert-based cross-validation, in just 40 epochs using 10-fold less non-augmented image data (Fig. 4d). It’s known that HeLa cells restrict *STm* growth by maintaining vacuole integrity; the small percentage of bacteria which escape vacuoles are decorated with ubiquitin and subsequently cleared by autophagy (23, 24). Interestingly, stage 2 HRMAn showed that the percent of cells which recruit ubiquitin to *STm* doubles upon IFNγ treatment, while the percent of decorated vacuoles/cell increases only slightly (Fig. 4e). As seen with *Tg* (Fig. 3e, S8a), IFNγ does not impact the distance of the ubiquitin coat to *STm* but increases its thickness (Fig. 4f). This indicates that more ubiquitin is recruited to cytosolic *STm* in the presence of IFNγ and growth of decorated bacteria was restricted (Fig. 4g). Consequently, although IFNγ treatment increases the number of cells that recruit ubiquitin to *STm* and the intensity of that recruitment, at the single-cell level HeLa cells appear to have reached their capacity for detection and autophagy-mediated clearance of cytosolic/ubiquitinated *STm* independent of IFNγ treatment (Fig. 4e-g).

**Fig. 4.**
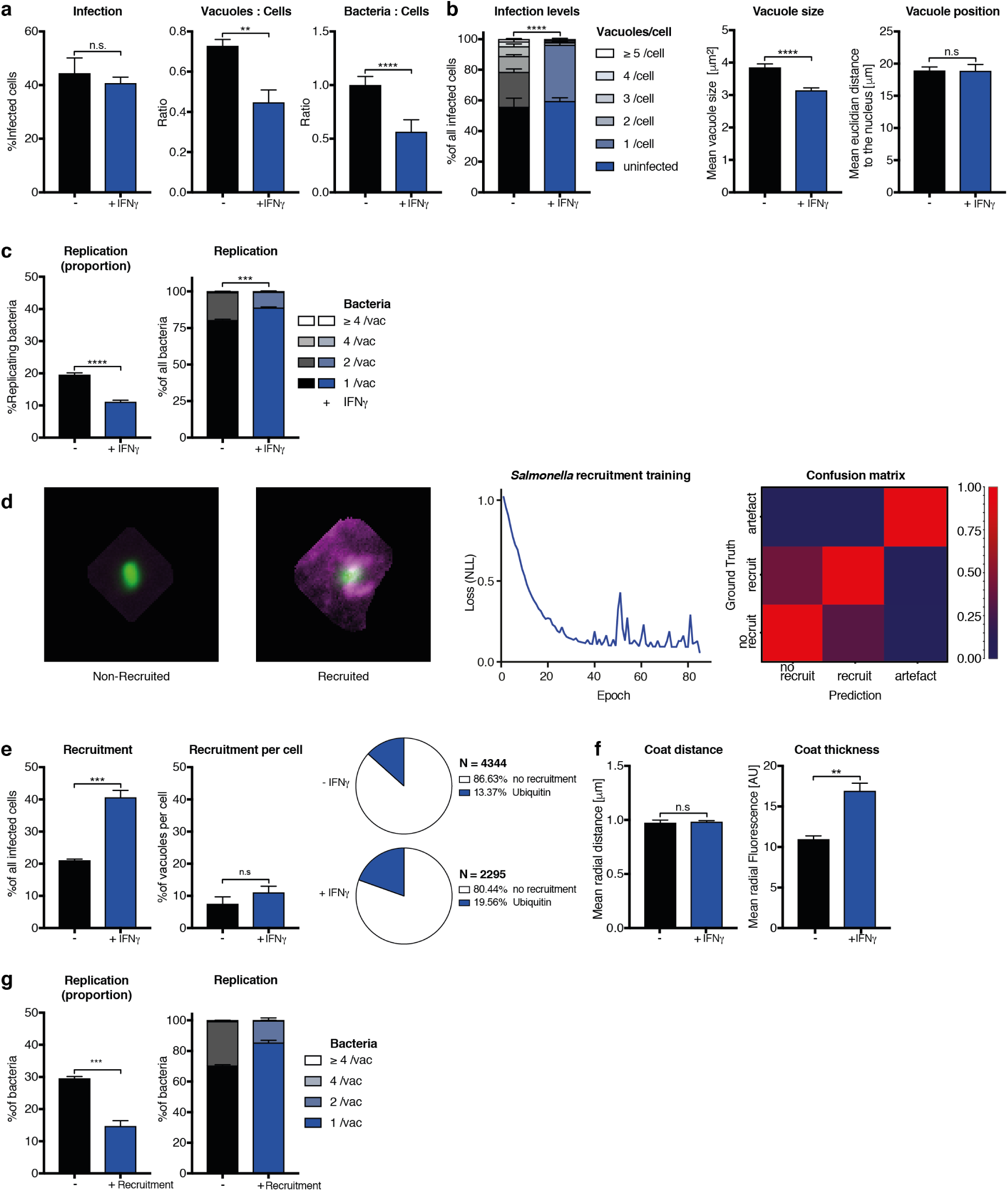
Analysis of Salmonella typhimurium infection in IFNγ-treated HeLa cells. HeLa cells were stimulated with 100 IU/mL IFNγ infected with *Salmonella typhimurium* (*STm*) and analyzed 2 hours post-infection. (a-c) Stage 1 infection analysis parameters. (a) Infection parameters depicted as total percent of *STm* infected cells, the ratio of *STm* vacuoles to cells and the ratio of bacteria to cells. (b) Cellular readouts showing the proportion of cells that contain a certain number of bacteria vacuoles, the mean vacuole size of *STm* and the vacuole position as the value of the mean euclidian distance of *STm* vacuoles to the host cell nucleus. (c) Replication capacity of *STm* shown as the proportion of replicating bacteria and the distribution of replicating *STm*. (d) Training of the deep convolution neural network (CNN) to analyze host protein recruitment to *STm* vacuoles and bacteria. Left: Example images showing the difference of no recruitment versus ubiquitin (magenta) recruitment to *STm*. Middle: Decrease of negative log likelihood (NLL) used as loss function during CNN training over training cycles (epochs) for *STm* model. Right: Confusion matrix of *STm* model validation, classification accuracy (0 to 1) during validation is color-coded blue to red. (e) Cellular response to infection with *STm* measured through the percentage of cells that decorate vacuoles and the average proportion of vacuoles per cell that are being decorated simultaneously and the overall proportion of ubiquitin decorated *STm* vacuoles. N shows the total number of vacuoles analyzed for each condition, percentages are indicated in the legend. (f) Properties of the host protein coat on *STm* vacuoles as the average coat distance for ubiquitin to *STm* and mean fluorescence intensity of ubiquitin. (g) Fate of *STm* grouped based on host protein recruitment. Shown is the proportion of replicating bacteria and the replication distribution based on recruitment status of the vacuole. All data shown above represents the mean of N = 3 experiments *±* SEM. Significance was determined using unpaired t-tests, n.s. = not significant, * p *≤* 0.0332, ** p *≤* 0.0021, *** p *≤* 0.0002 and **** p *<* 0.0001

## Discussion

Recent advances have made deep CNNs a powerful image analysis method (18, 25). Inspired by abstraction of animal visual cortex architecture, CNNs are able to generalize patterns independent of minor phenotypic differences (26, 27). Combining automated image segmentation, machine learning and a deep CNN in an ensemble, HRMAn is a powerful open-source, user-friendly software for the analysis of host-pathogen interaction at the single-cell level. To date, HRMAn represents the only open-source CNN-driven host-pathogen analysis solution for fluorescent images. Many automated image analysis programs, some of which incorporate machine learning elements, have been developed and are successfully used for classical image segmentation (Table 1). However, when presented with the problem of classifying host protein recruitment to a pathogen, inaccurate classical image segmentation could lead to erroneous results. Employing an intuitive open-source artificial intelligence algorithm, HRMAn circumvents these problems and delivers user-defined automated and unbiased enumeration of this subset of the host-pathogen interplay. Using *Tg* and *STm* infection models we demonstrate that HRMAn is capable of detecting and quantifying multiple pathogen and host parameters. Designed for biologists, HRMAn requires no coding or specialized computer science knowledge. Its modular architecture and the use of KNIME providing a graphical representation of the analysis pipeline, allows users to tailor experimental outputs to their own datasets and questions. As the models we have generated can be used as primers to lower the training dataset size, computation power and training time requirements, HRMAn can be rapidly applied to similar large-scale, image-based experimental setups. As such, HRMAn will allow a broad range of researchers to extend into the realm of high-throughput, unbiased, quantitative, single-cell analysis of host-pathogen interaction.

## Author Contribution

DHF and EMF conceived the idea for HRMAn, DHF and AY designed and implemented HRMAn, DHF performed experiments, BC provided essential experimental protocols, BC, JW, MB tested HRMAn in multiple settings, MH provided essential high content imaging guidance and performed the automated image acquisition. DHF, AY, JPM and EMF wrote the manuscript. All authors contributed to analysis and interpretation of the data and revision of the manuscript.

## Competing Financial Interests

The authors declare no competing financial interest.

## ACKNOWLEDGEMENTS

We thank all members of the Frickel lab for productive discussion. This work was supported by the Francis Crick Institute, which receives its core funding from Cancer Research UK (FC001076), the UK Medical Research Council (FC001076), and the Wellcome Trust (FC001076). EMF was supported by a Wellcome Trust Career Development Fellowship (091664/B/10/Z). DHF was supported by a Boehringer Ingelheim PhD fellowship. AY, JM were supported by core funding to the MRC Laboratory for Molecular Cell Biology at University College London (J.M.), the European Research Council (649101-UbiProPox), the UK Medical Research Council (MC_UU12018/7).

## Methods

### Code availability

All open-source KNIME workflows used in this publication can be found at: http://bit.ly/hrman2018. Upon publication HRMAn framework will be deposited on GitHub and the homepage http://hrman.org under GPLv3 open-source software license to allow for rapid and open dissemination and free availability for the research community. The models and their respective weights obtained through training will be deposited on GitHub and the homepage as well.

### Image acquisition

For simple infection analysis (stage 1), 96-well plates (see Microscopy sample generation) were imaged on an ArrayScan™ VtI Live High Content Imaging Platform (Thermo Scientific) using 20x magnification and depending on the experiment, 15-20 fields of view per well. For recruitment analysis to *Toxoplasma gondii (Tg)* vacuoles, glass-bottom 96-well plates were imaged on an ArrayScan™ VtI Live High Content Imaging Platform but using 40x magnification and depending on the experiment, 50 fields of view per well. In both cases, following image acquisition, the images were exported from HCS Studio Cell Analysis Software as single channel 16-bit tiff files before they were fed into the HRMAn analysis pipeline.

For recruitment analysis to *Salmonella typhimurium (STm)* vacuoles, images of coverslips were acquired on a Ti-E Nikon microscope equipped with an LED-illumination and an Orca-Flash4 camera using a 60x magnification. 75 fields of view per coverslip were acquired using multi-position acquisition. Images were exported as single channel 16-bit tiff files with Nikon NIS Elements software before they were fed into the HRMAn analysis pipeline.

Generally, HRMAn can work with any common image file format, but the use of uncompressed, lossless formats like tiff (or png) is recommended. Furthermore, HRMAn can work with images acquired on any type of fluorescence microscope and is truly platform independent.

### Image analysis using HRMAn

Following image acquisition, the images were loaded into the HRMAn pipeline. Images can be in any common file format, preferably as single-channel tiff files. The used image reader loads images from all file formats supported by the Bio-Formats library (a list of the supported formats can be found here: http://loci.wisc.edu/bio-formats/formats). If the images were not acquired on a high-content imaging platform, they can be renamed with HRMAn to mimic the file names and the plate format. This is needed to cluster the output data and perform error calculation. Furthermore, the OME-XML-metadata is loaded and information on the image is extracted (e.g. image size, type and origin). While the images are loaded into KNIME, the user is asked to provide some basic information on the image acquisition and the type of analysis to be performed. This includes the used magnification, type of analysis, channel number and order and providing a plate map to cluster the data.

HRMAn then pre-processes the images, meta-data and provided information and lets the user inspect the input images arranged into a grid and sorted by the field of view. Next the input images undergo illumination correction by dividing the background as a mean image of all acquired images in a channel-wise fashion. Following this step, the individual channels are segmented to detect the Nuclei, the pathogens and the cells:

(1) Nuclei are detected using Otsu’s method thresholding, a watershed and connected component analysis. Fields of view containing insufficient numbers of nuclei (i.e. empty fields) are excluded from the following analysis.
(2) The pathogens (or vacuoles) are detected after image normalization and filtering through thresholding using Otsu’s method. Incomplete labels are corrected by filling holes and pathogen vacuoles are separated through watershedding. Labels are created with a connected component analysis.
(3) Cell labels are created using Huang thresholding and a Voronoi segmentation using the nuclei labels as starting points. Optionally the images can be enhanced using Contrast Limited Adaptive histogram equalization (CLAHE) to improve the segmentation accuracy. All cell labels touching the border of an image are excluded from the analysis. Furthermore, the created labels are filtered based on their size and the user-defined parameters such as magnification and detector size. The filter values for *STm* and *Tg* were empirically determined using thousands of images from different experiments. Based on which pathogen type is chosen by the user, HRMAn will adjust the filters automatically. Using labelling arithmetic, pathogen labels that are not contained within a cell label are removed from the dataset, as they represent extracellular pathogens. The created labels can then be inspected by the user through an interactive label viewer displaying the original image next to the labels.

Using the created labels, the infection readouts for stage 1 are created: First, cell numbers *N*_*Cells*_ and vacuole numbers *N*_*V* *acuoles*_ are determined by counting the numbers of respective labels in all acquired fields per well (= replicate). Using these values, the vacuole to cell ratio is calculated:

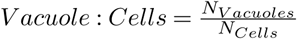

Next, the dependencies between the cell labels and vacuole labels are used to calculate the proportion of infected cells and the infection levels of the cells. The label dependencies determine which vacuole labels *V*_1_, *V*_2_, …, *V*_*i*_ are contained by which cell label *C*_1_, *C*_2_ …, *C*_*j*_. If a cell label *C* contains at least one vacuole label *V*, the cell is considered as infected cell *C*_*inf*_. This is used to calculate the proportion of infected cells:

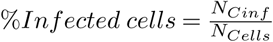

with *N*_*Cinf*_ as the number of infected cells *C*_*inf*_

To determine the more precise distribution describing how many vacuoles are contained by which proportion of cells (= Infection levels) the cells *C* are split into subgroups according to the number of vacuoles they contain (no vacuoles or uninfected = *C*_0_, 1 vacuole = *C*_1_, …, 5 or more vacuoles per cell = *C*_≥5_) and then the proportion is calculated:

0 vacuoles per cell (uninfected):

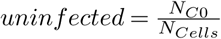

1 vacuole per cell:

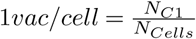

…

5 or more pathogens per vacuole:

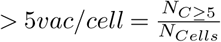

Also based on the dependencies, the mean euclidean distance d between the centroid of the vacuole labels (with coordinates *X*_*V*_ and *Y*_*V*_) within a cell and its nucleus’ centroid (with coordinates *X*_*N*_ and *Y*_*N*_) is determined as the position of the vacuole:

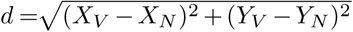

Using the vacuole labels, of vacuoles contained within cells, and working on the original images the properties of the vacuoles are measured as mean values for each well. These include mean vacuole size, shape descriptors (Circularity, Perimeter, Convexity, Extent, Diameter) and fluorescence properties (Minimum, Mean and Maximum Fluorescence). Using the above determined values as attributes, a decisiontree machine learning algorithm determines good classifiers and employs them to classify each vacuole label Vi based on how many individual pathogens *P*_*V* *i*_ it contains. This step requires providing an annotated dataset.

Based on this classification the vacuoles can be divided into individual groups for the number of pathogens they contain (1*/vac* = *V*_1*V* *ac*_, 2*/vac* = *V*_2*V* *ac*_, 4*/vac* = *V*_4*V* *ac*_ and 4*ormore/vac* = *V* _≥4*V* *ac*_) and the number of vacuoles in each group is counted (e.g. number of vacuoles that contain just one pathogen = *N*_*V* 1*V* *ac*_). To calculate the proportion of replicating pathogens, the number of vacuoles that contain at least two pathogens is divided by the total number of vacuoles:

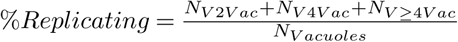

Similarly, the individual proportions of the vacuole groups are calculated to illustrate pathogen replication distribution:

1 pathogen per vacuole:

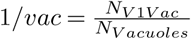

2 pathogen per vacuole:

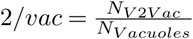

4 or more pathogens per vacuole:

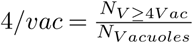

Combining the information on the number of vacuoles and the number of pathogens *P*_*V*_*i* each individual vacuole *V*_*i*_ contains, the total number of pathogens *N*_*P* *athogens*_ is calculated:

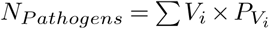

This can be used to determine the Pathogen load by normalization to the cell number:

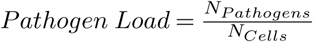

This concludes stage 1 infection analysis performed by HRMAn. In the end of the analysis, the values calculated for each well or replicate is combined with the values for the other wells belonging to the same sample group based on the user-provided plate map and error calculation is performed. If the user decides to perform only stage 1 infection analysis the HRMAn image analysis pipeline will stop here, if host protein recruitment analysis is to be performed the data will be fed into the second stage for which the implemented deep Convolutional Neural Network (CNN) has to be trained first.

### Deep Learning Setup and Neural Network Architecture

To classify pathogen recruitment, we employed a deep Convolutional Neural Network (CNN) adopted from a published AlexNet (10, 28) (Fig. S1). To ensure our neural network can be implemented by other researchers with no coding, it was based on the open source DeepLearning4J library implementation in Eclipse-KNIME. Unlike the original AlexNet our network was suited to take 100 by 100 by 2 pixels images as input and designed to run on a single graphic processing unit (GPU). Since our fluorescence microscopy input data had two fluorescence channels, rather than the standard RGB dimension of three, we used two to save computational resources. Furthermore, the choice of DeepLearning4J as a deep learning library allowed us to use 16-bit microscopy images directly, preventing information loss upon conversion of scientific imaging data. Our deep learning hardware was based on a single Nvidia 1080 Ti GPU set up in Intel®Core™ i7 4790K system equipped with 32 Gb of RAM and a SSD. Our architecture consisted of a total of 5 convolutional layers, where the first two were immediately followed by local response normalization layers and max pooling layers and the last three convolutional layers were followed by one max pooling layer connected to a fully connected layer. All these layers used rectified linear unit (ReLU) as activation function (29). Finally, to accomplish the classification task the network had a fully connected layer with a soft maximum (SoftMax) activation (30).

### Neural Network Training and Hyperparameters Optimization

Our neural networks were trained using stochastic gradient descent (SGD) based backpropagation on the augmented original data over at least 80 cycles (epochs) (31, 32). To fully utilize the GPU capacity, training was performed in mini-batches of 200. To improve the SGD performance, we utilized an ADAM updater with ADAM Mean Decay of 0.9 and ADAM Variance Decay of 0.999 (33). We employed the Xavier algorithm for the weight initialization strategy and negative log likelihood (NLL) as our loss function (33). We used learning rates between 0.001 and 0.01 adjusted accordingly to ensure the optimal loss curve decay during training. Together with the weights initialization strategy and the updater choice these were the main hyperparameters optimized in multiple iterations to ensure good training process.

### Data Preparation, Augmentation and Model Validation

Vacuole images used for creation of our deep learning model were segmented from large field of view micrographs obtained from high-content imaging. To ensure the dimension of the images are uniform, we padded all vacuole with zero-value padding to a uniform 100 by 100 pixels size. Next, we manually labeled the segmented vacuoles into recruited, non-recruited and artefactual (in case of erroneous segmentation of the vacuole). This labeled dataset was then split into the training and test datasets. To ensure our neural network has sufficiently diverse learning data, upon splitting the original labeled dataset into training and test subsets we performed data augmentation using a custom developed macro for ImageJ. During the augmentation processing, the original labeled dataset was concatenated with a modified version of it. The modifications included various rotations, image reflecting, image translation within the field of view. As microscopy data is typically rotation-, translation-or reflection-invariant, such modification allowed us to create a better dataset aiming at a more generalized model. Model validation was performed using the non-augmented test fraction of the labeled dataset previously unseen by the model. For this, we used the trained model as first input and passed the labeled test data through the classifier in the second input. The classification accuracy was assessed by accuracy score, numbers of true positive, false positive, true negative and false negative, as well as Cohen’s kappa values. A direct summary of the accuracy was visualized in a confusion matrix illustrating a mismatch between original label (Ground Truth) and the class assigned by the classifier (Fig. 2).

### Host protein recruitment analysis in HRMAn

For recruitment analysis, the vacuole labels created in stage 1 of the analysis are dilated over 20 iterations to create non-overlapping regions of interest (ROIs) around them. Simultaneously, the fluorescence images of the pathogen and the respective channel with fluorescence signal of the host protein are merged into a dual channel image. The created ROIs are used to crop the dual channel images, which creates images of the pathogen and its surrounding stained host protein. The images are clipped to 100 by 100 pixel and fed into a feedforward predictor (classification) which uses the provided and trained deep convolution neural network (CNN) to classify the pathogens vacuoles based on their coating. Once the vacuoles are separated into two groups, they are analyzed with the above described methods of stage 1 infection analysis but additionally comparing recruited versus non-recruited vacuoles. Thus, in addition to the overall infection parameters from stage 1 the user is provided with the same parameters but further layered for the cellular response. In the case of co-recruitment analysis, two images of each vacuole are created with both containing the pathogen signal, but each containing a different second channel, representing the different stainings. After classification with the CNN, the vacuoles can be compared for recruitment or co-recruitment and all the above described parameters are calculated for them individually. Using the previously determined label dependencies of vacuoles *V*_*i*_ and cells *C*_*i*_ and the classification of the vacuoles *V*_*i*_ by the CNN, HRMAn can furthermore calculate the proportion of cells that do respond to infection by decorating at least one vacuole and the proportion of vacuoles decorated per cell, if a single cell contains more than one pathogen vacuole. Furthermore, working only on the decorated vacuoles, we used a custom-made Fiji code to create a pixel-wise radial intensity profile starting from the pathogen centroid. The distance of the maximum fluorescence intensity is then used to define the distance of the coat from the pathogen center. Moreover, the mean fluorescence intensity of the coat is determined and can be used as a readout for the amount of protein recruited to each pathogen vacuole. Finally, all mean values and errors for the replicate conditions, as defined by the user’s plate map, are calculated and written into a single spreadsheet file. Before this, the user can also define a scaling factor between pixel and actual metric values which will adjust the output values from pixel (px) to µm or to µm2 respectively.

### Cell culture

THP-1 (ATCC) were maintained in RPMI with GlutaMAX (Life Technologies) supplemented with 10% FBS (Sigma), at 37°C in 5% CO_2_. THP-1s were differentiated with 50 ng/mL phorbol 12-myristate 13-acetate (PMA) for 3 days and then rested for 2 days by replacing the differentiation medium with complete medium without PMA. Cells were not used beyond passage 20. Human Umbilical Vein Endothelial cells, HUVECs, (Promocell C12203), were maintained in M199 medium (Life Technologies) supplemented with 30 g/mL endothelial cell growth supplement (ECGS) (Upstate 02–102), 10 units/mL heparin (Sigma H-3149) and 20% FBS (Sigma). Cells were grown on plates, pre-coated with 1% (w/v) porcine gelatin (Sigma G1890) and cultured at 37°C in 5% CO_2_. HUVEC were not used beyond passage 6. HeLa (ECACC, Sigma), A549 (ATCC) and human foreskin fibroblasts, HFFs (ATCC), were cultured in DMEM with GlutaMAX (Life Technologies) supplemented with 10% FBS (Sigma), at 37°C in 5% CO_2_. HeLa and A549 cells were not used beyond passage 25 and HFFs were not used beyond passage 15. All cell culture was performed without addition of antibiotics and the cells were regularly tested for mycoplasma contamination by immunofluorescence, PCR and agar test.

### Interferon stimulation of cells

All five cell lines used in this publication were stimulated for 16 h in complete medium at 37°C with addition of 100 IU/mL human IFNγ (R&D Systems) prior to infection, if not indicated otherwise.

### *Toxoplasma gondii* infection

Parasites were always passaged the day before infection onto new HFFs to obtain parasites with a high viability for infection. *Tg* were prepared from freshly 25G syringe-lysed HFF cultures in 10% FBS by centrifugation at 50 x g for 3 minutes and transferring the cleared supernatant into a new tube and subsequent centrifugation at 500 x g for 7 minutes and re-suspension of the pelleted parasites into fresh complete medium. Then, the parasites were added to the experimental cells at a MOI of 3 for both type I and type II strains. The cell cultures with added *Tg* were then centrifuged at 500 x g for 5 minutes to synchronize the infection. Two hours post-infection, the cultures were thoroughly washed two times with warm PBS to remove any uninvaded parasites and fresh complete medium was added prior to culturing at 37°C, 5% CO_2_ for the required time.

### Bacteria culture and infection

*Salmonella enterica serovar Typhimurium* 12023 wild-type strain containing the plasmid pFVP25.1, carrying gfpmut3A under the control of the rpsM constitutive promoter were grown in Luria Bertani (LB) medium supplemented with ampicillin (50 µg/ml). Prior to infection, bacteria were grown to induce SPI-1 T3SS expression: cultures of *STm* were grown at 37°C in LB, diluted 1:50 into fresh LB containing 300 mM NaCl the next morning and incubated shaking at 200 rpm until OD600 = 0.9 – 1.0 was reached. Bacteria were washed in medium without FBS before use. Cells were infected at a MOI of 50 and infections were synchronized by centrifuging bacteria onto the cells at 750 x g for 10 minutes. 15 minutes post infection, the cells were thoroughly washed three times with warm PBS to remove extracellular bacteria and medium containing 100 µg/mL Gentamicin (Gibco) was added for 30 min. Then, Gentamicin concentration was reduced to 10 g/mL and cells were incubated further at 37°C, 5% CO_2_ for the appropriate amount of time.

### Antibodies

Antibodies used in this study were rabbit pAb anti-p62 (MBL, #PM045), mouse mAb anti-GRA2 (Biovision, A1298) and mouse mAb anti-ubiquitin (FK2) (Enzo Lifesciences, PW8810). Secondary antibodies used were Alexa Fluor 647-conjugated goat anti-rabbit or anti-mouse and Alexa Fluor 568-conjugated goat anti-mouse (Molecular Probes).

### Microscopy sample generation

#### Simple infection analysis

For simple infection analysis, 30,000 THP-1s per well were seeded 5 days prior to IFNγ treatment and differentiated with 50 ng/mL PMA for three days and then rested for 2 days in complete medium. HFFs were harvested by washing a confluent monolayer with PBS and subsequent lifting of the cells with 0.05% Trypsin-EDTA (Gibco). Cells were centrifuged at 250 x g for 5 mins, re-suspended in fresh DMEM plus 10% FBS and 20,000 HFFs per well were seeded the day before IFNγ treatment. Similarly, HUVECs were harvested and 15,000 cells per well were seeded in complete medium the day before IFNγ treatment. A549s and HeLa cells were harvested in the same way and 8,000 cells per well were seeded the morning before IFNγ treatment. All cells were seeded on 1% (w/v) porcine gelatin pre-coated black-wall, clear bottom 96-well plates (Thermo Scientific). In the evening, all cells were treated with 100 IU/mL IFNγ or medium and left at 37°C, 5% CO_2_ overnight. The next morning the cells were infected with either *Tg* or *STm* as described above. After the appropriate infection duration the infected cells were again thoroughly washed with warm PBS to remove as many uninvaded pathogens as possible and subsequently fixed with 4% methanol-free paraformaldehyde (Thermo Scientific). Fixed specimens were permeabilized with PermQuench buffer (0.2% (w/v) BSA and 0.02% (w/v) saponin in PBS) for 30 minutes at room temperature. Then PermQuench buffer containing 1 g/mL Hoechst 33342 (Life Technologies) and 2 ug/mL CellMask™ Deep Red plasma membrane stain (Invitrogen) were added and samples were incubated at room temperature for 1 hour. After staining, the specimens were washed with PBS 5 times and kept in 200 L PBS per well for imaging.

#### Recruitment analysis

For recruitment analysis, the cells were prepared as described above, but they were seeded on 1% (w/v) porcine gelatin pre-coated black-wall, glass bottom 96-well imaging plates CG 1.0 (Miltenyi Biotec) to allow higher resolution imaging. After fixation, cells were permeabilized identically and then stained with primary antibody diluted in PermQuench buffer for 1 hour at room temperature. After three washes with PBS, cells were incubated with the appropriated secondary antibody and 1 µg/mL Hoechst 33342 diluted in PermQuench buffer for another hour at room temperature. Then, the specimens were washed with PBS 5 times and kept in 200 µL PBS per well for imaging. In the case of recruitment analysis to *STm* vacuoles, the cells were seeded on 1% (w/v) porcine gelatin pre-coated 9 mm coverslips in 24-well plates. After fixation and identical staining procedure, the coverslips were mounted using 5 L ProLong™ Gold Antifade Mountant (Invitrogen).

### Data handling and statistical measurements

Data was plotted using GraphPad Prism and presented with error bars as standard error of the mean (SEM). Significance of results was determined by non-parametric one-way ANOVA, two-way ANOVA or unpaired t-test as indicated in the figure legends.

## Supplementary Information

This is supplementary information to the preprint manuscript from *Fisch & Yakimovich et al. 2018*. Supplementary section contains Supplementary table, supplementary figures and figure legends. All further information is available upon request.

## Supplementary Figures

**Fig. S1:**
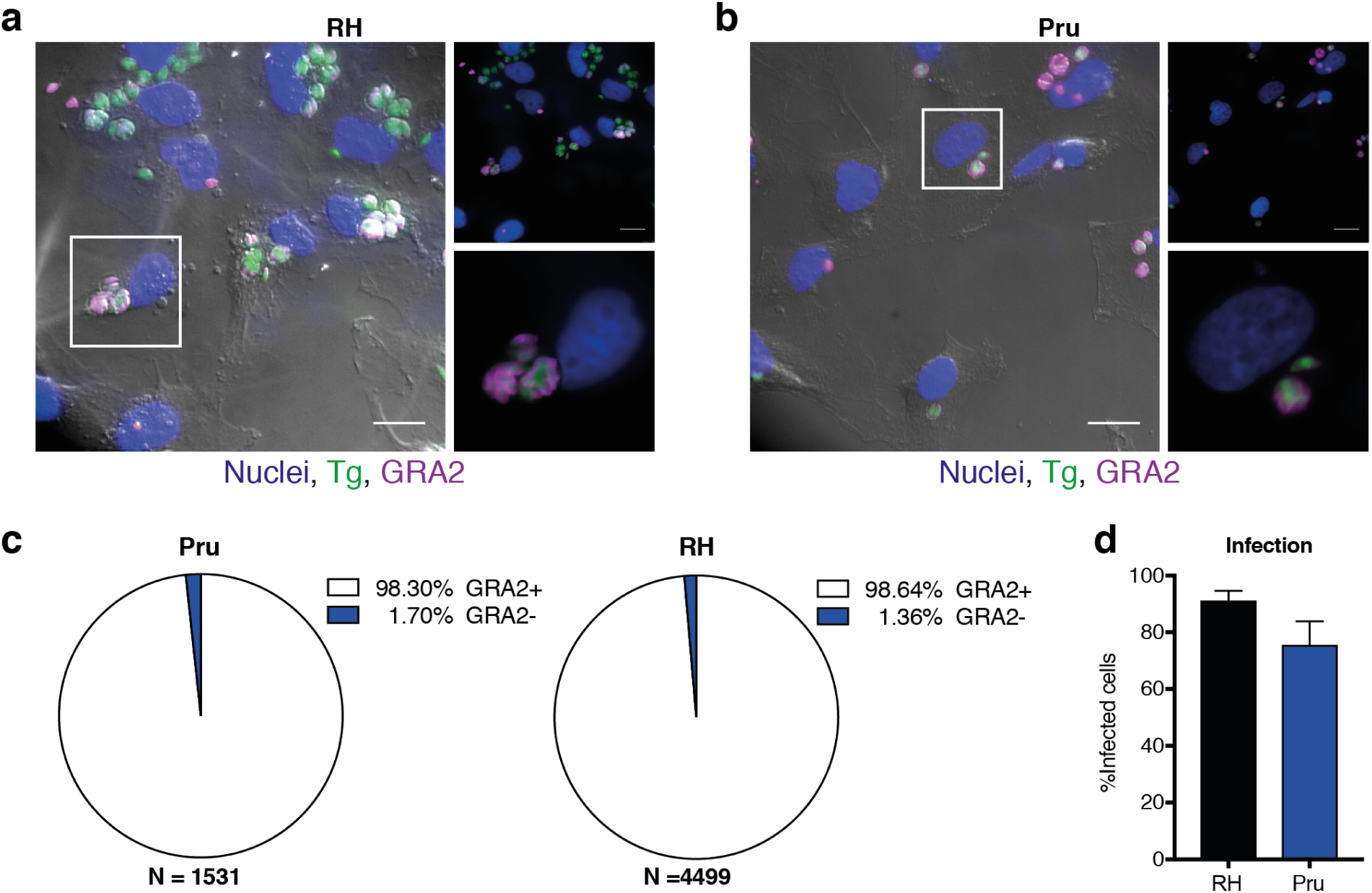
Infection of HeLa cells with *Toxoplasma gondii* at 6 hours post-infection. (a+b) HeLa cells were infected with either type I (RH) Toxoplasma gondii (a) or type II Pru *Toxoplasma gondii* and underwent a stringent washing procedure to eliminate uninvaded parasites. Infected cells were stained with anti-GRA2 (purple) to illustrate vacuole establishment. Scale bar indicates a distance of 20 m. (c) Quantification of GRA2 positive vacuoles for type I and type II *Toxoplasma gondii* infected cells. (d) Quantification of infected cells as proportion of all captured cells. Data shown in (c) and (d) represents the mean of N = 3 experiments ±SEM, N = total number of vacuoles analyzed in the course of three experiments.

**Fig. S2:**
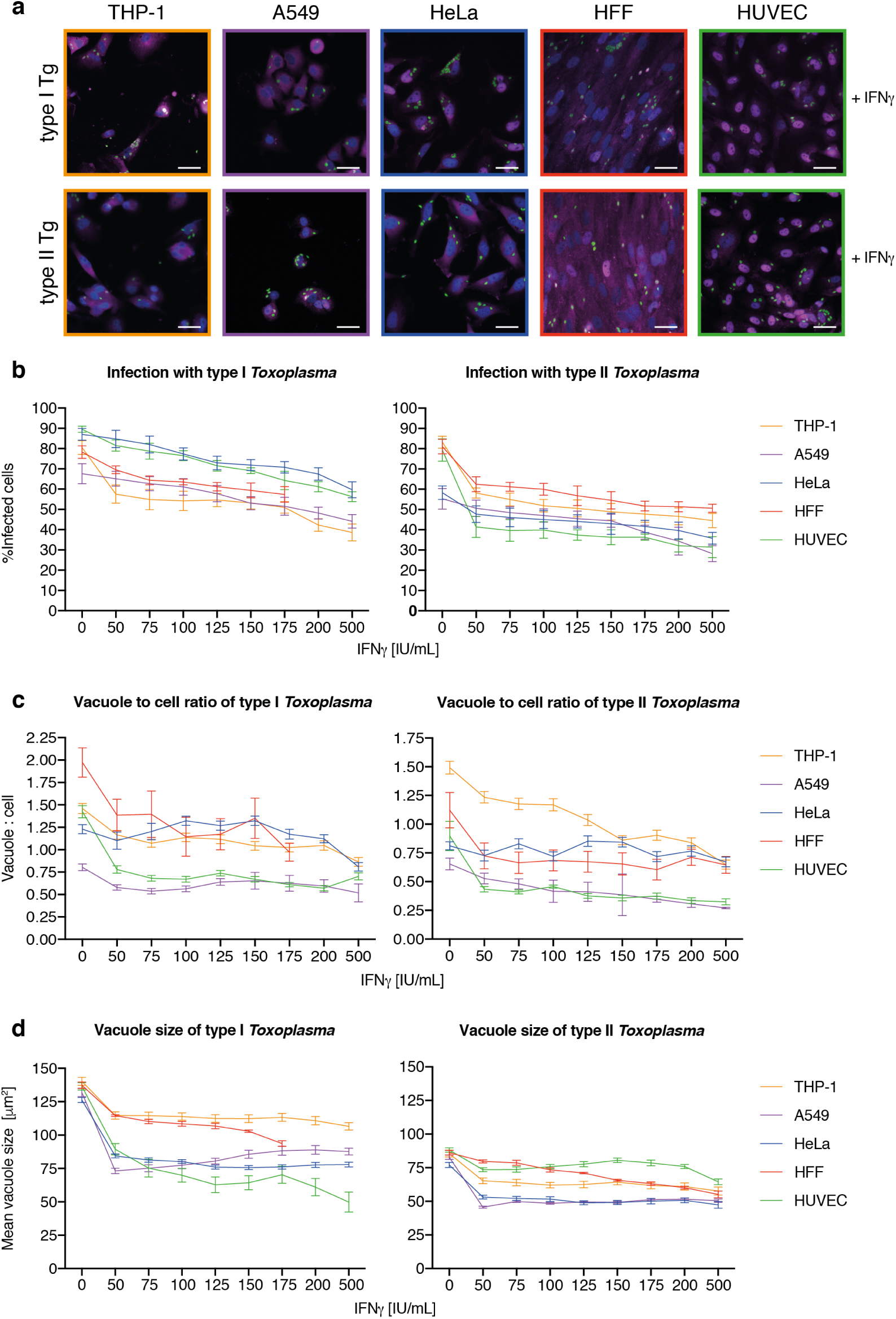
IFNγ dose-dependent killing and replication-inhibition of *Toxoplasma gondii* in 5 human cell types at 24h postinfection. (a) Example images employed to analyze the cellular response to *Toxoplasma gondii (Tg)* infection dependent on different dosages of IFNγ pre-treatment for these 5 different human cell lines: macrophage-like, PMA-differentiated THP-1s (yellow), alveolar-epithelial tumor cells A549 (purple), the cervical cancer cell line HeLa (blue), human foreskin fibroblasts (HFF, red) and primary human umbilical vein endothelial cells (HUVEC, green). Scale bar indicates a distance of 30 m. All images represent conditions pre-treated with 100 IU/mL IFNγ. (b-d) Host-pathogen interaction parameters of *Tg* type I and II infection were analyzed with HRMAn 24 hours post-infection. (b) Percent *Tg* infected cells, (c) ratio between *Tg* vacuoles and cells and (d) the mean vacuole size of *Tg*. All data shown above represents the mean of N = 3 experiments ±SEM.

**Fig. S3:**
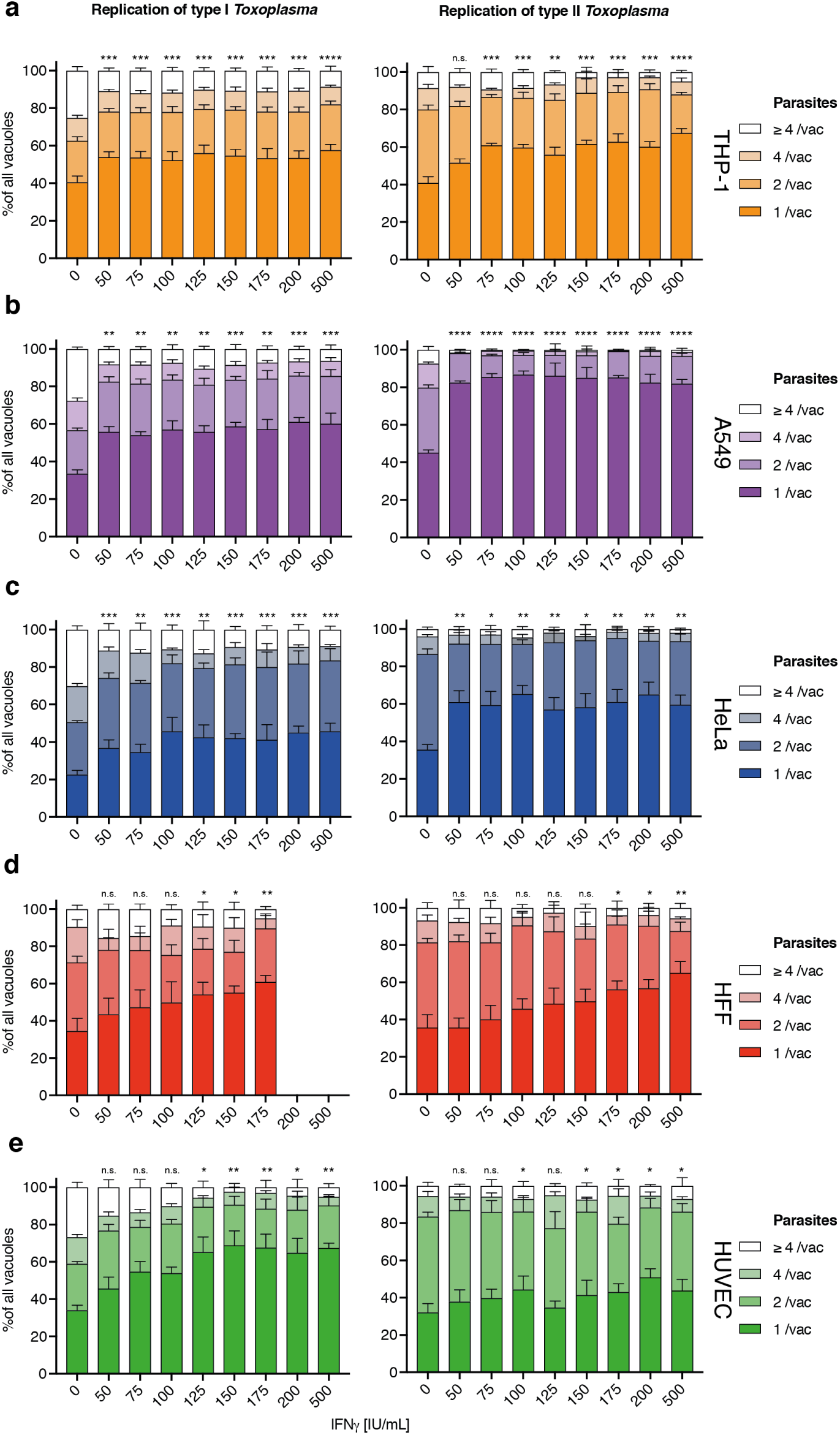
IFNγ dose-dependent replication-inhibition of *Toxoplasma gondii* in 5 human cell types analyzed as parasites per vacuole at 24h post-infection. (a-e) Mean vacuole size of *Toxoplasma gondii (Tg)* dependent on different dosages of IFNγ pre-treatment for 5 different human cell lines converted to number of parasites per vacuole as per HRMAn decision tree machine learned algorithm. Plotted are the distribution of vacuoles that contain one parasite, two, four or more than 4 parasites. Data shown was recorded 24 hours post-infection. Growth restriction of type I (RH) *Tg* (left) or type II (Pru) *Tg* (right) in THP-1 cells (a), A549 cells (b), HeLa cells (c), HFF (d) and HUVECs (e). All data shown above represents the mean of N = 3 experiments ±SEM. Significance was determined using non-parametric one-way Anova, n.s. = not significant, * p *≤* 0.0332, ** p *≤* 0.0021, *** p *≤* 0.0002 and **** p *<* 0.0001.

**Fig. S4:**
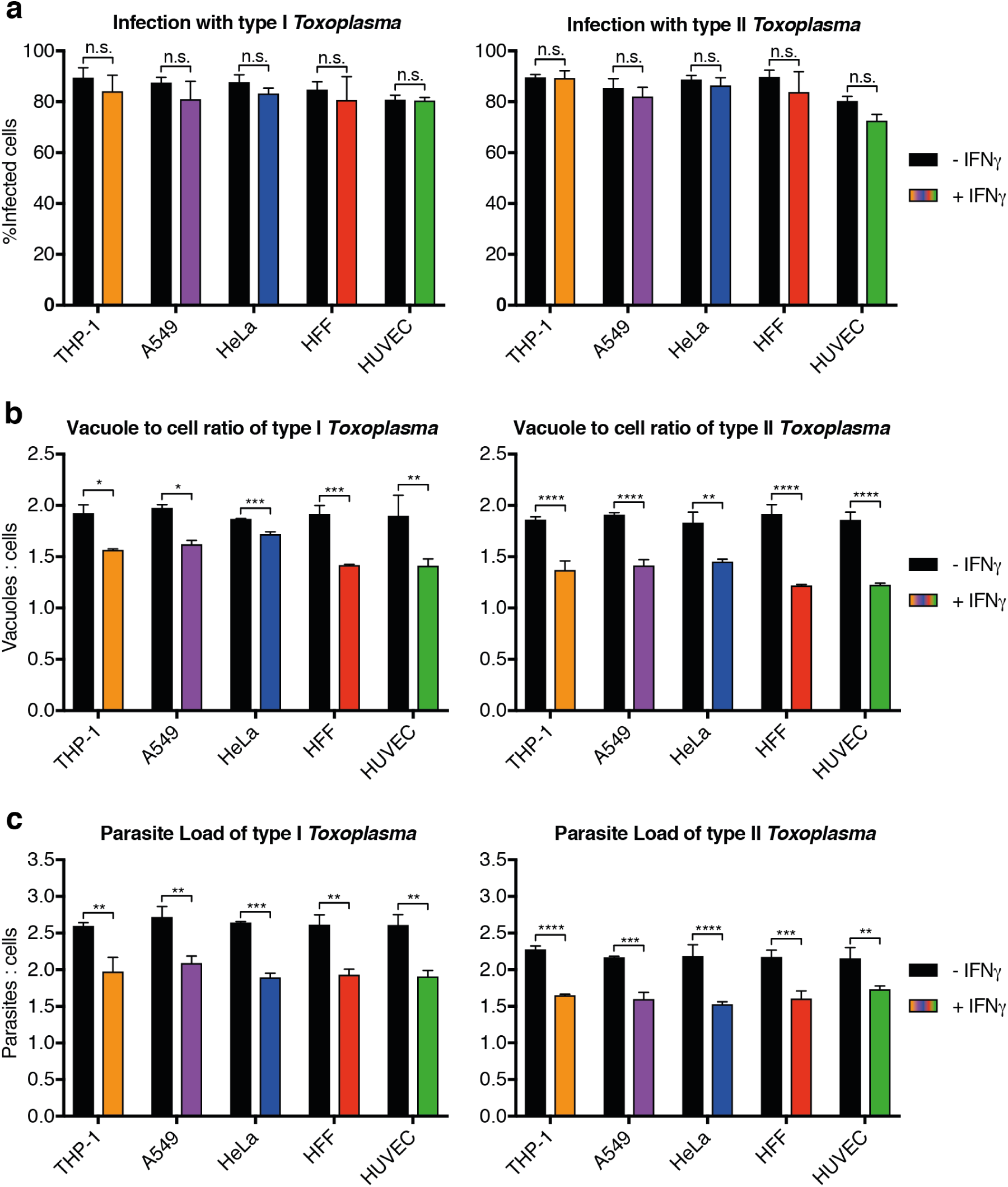
Systematic analysis of IFNγ-dependent cellular control of *Toxoplasma gondii* infection of 5 human cell types at 6h post-infection. Analysis of the proportion of cells infected with type I (RH) and type II (Pru) *Toxoplasma gondii (Tg)* in IFNγ-treated 5 human cell types. (a) Total percent infected cells for all cell lines tested, (b) the ratio of *Tg* vacuoles to cells and (c) the ratio of total number of individual parasites to cells. All data shown above represents the mean of N = 3 experiments ±SEM. Significance was determined using unpaired t-tests, n.s. = not significant, * p ≤0.0332, ** p ≤0.0021, *** p ≤0.0002 and **** p *<* 0.0001.

**Fig. S5:**
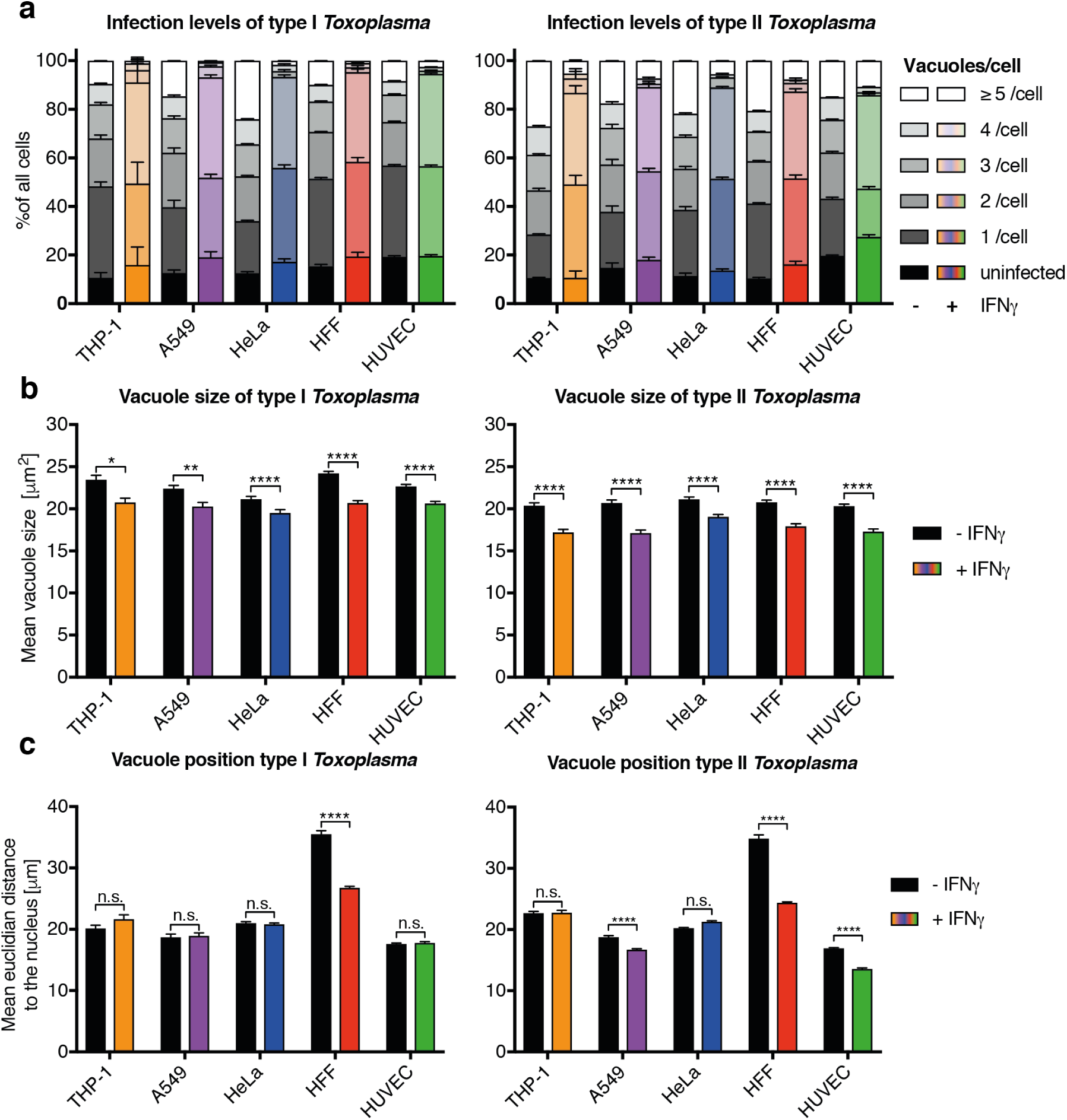
Systematic analysis of IFNγ-dependent replication control of *Toxoplasma gondii* infection of 5 human cell types at 6h post-infection. Measuring the infectivity and position of type I (RH) and type II (Pru) *Toxoplasma gondii (Tg)*. (a) The proportion of cells that contain a certain number of parasite vacuoles, (b) the mean vacuole size of *Tg*, (c) Value of the mean euclidian distance of *Tg* vacuoles to the host cell nucleus. All data shown above represents the means of N = 3 experiments ±SEM. Significance was determined using unpaired t-tests, n.s. = not significant, * p≤ 0.0332, ** p≤ 0.0021, *** p ≤0.0002 and **** p *<* 0.0001.

**Fig. S6:**
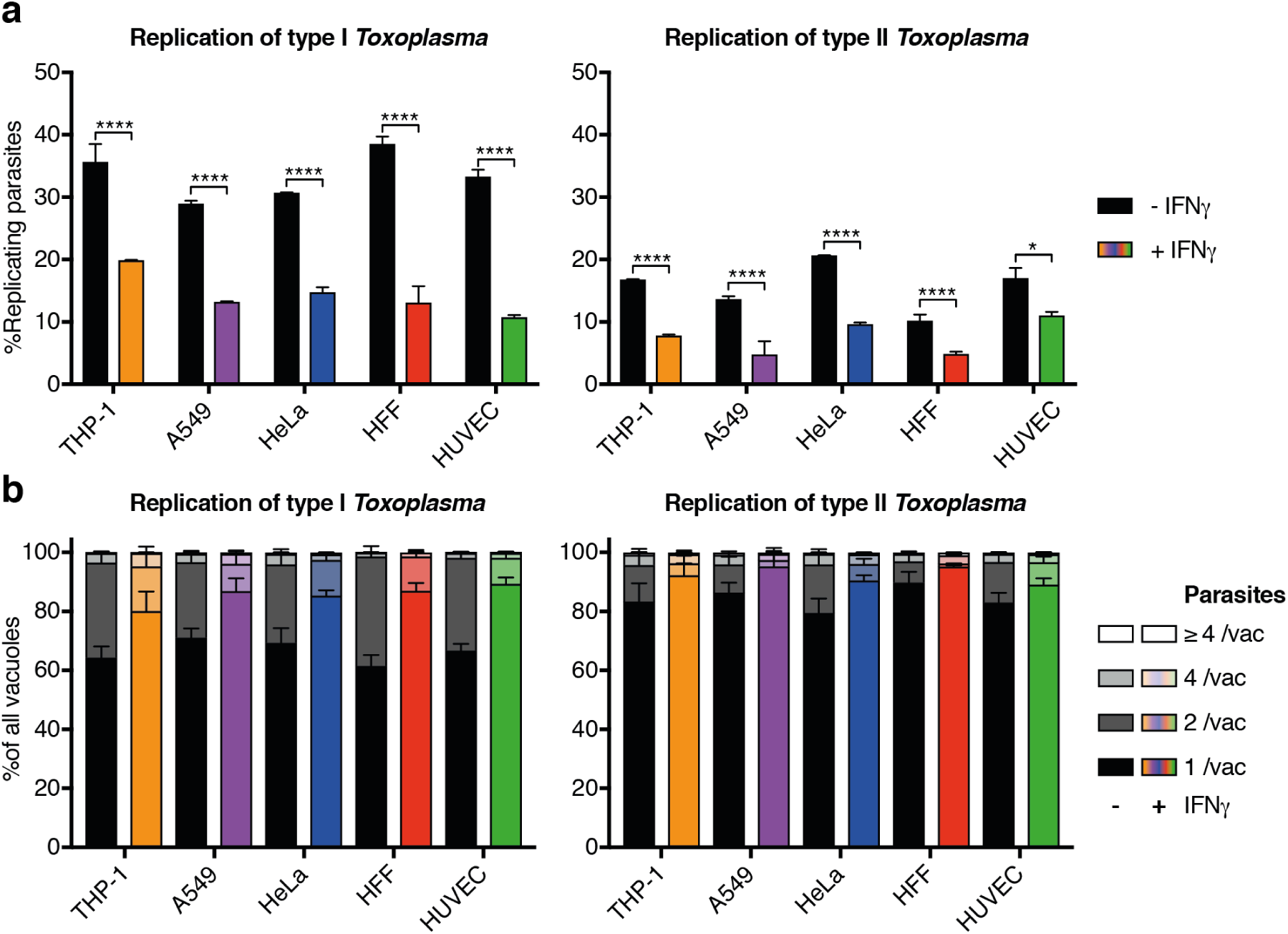
Systematic analysis of IFNγ-dependent replication control of *Toxoplasma gondii* infection of 5 human cell types at 6h post-infection analyzed as parasites per vacuole. Measuring the replication capacity of type I (RH) and type II (Pru) *Toxoplasma gondii (Tg)*. (a) The proportion of replicating parasites, (b) the distribution of replicating *Tg*. All data shown above represents the means of N = 3 experiments ±SEM. Significance was determined using unpaired t-tests, n.s. = not significant, * p *≤* 0.0332, ** p *≤* 0.0021, *** p *≤* 0.0002 and **** p *<* 0.0001.

**Fig. S7:**
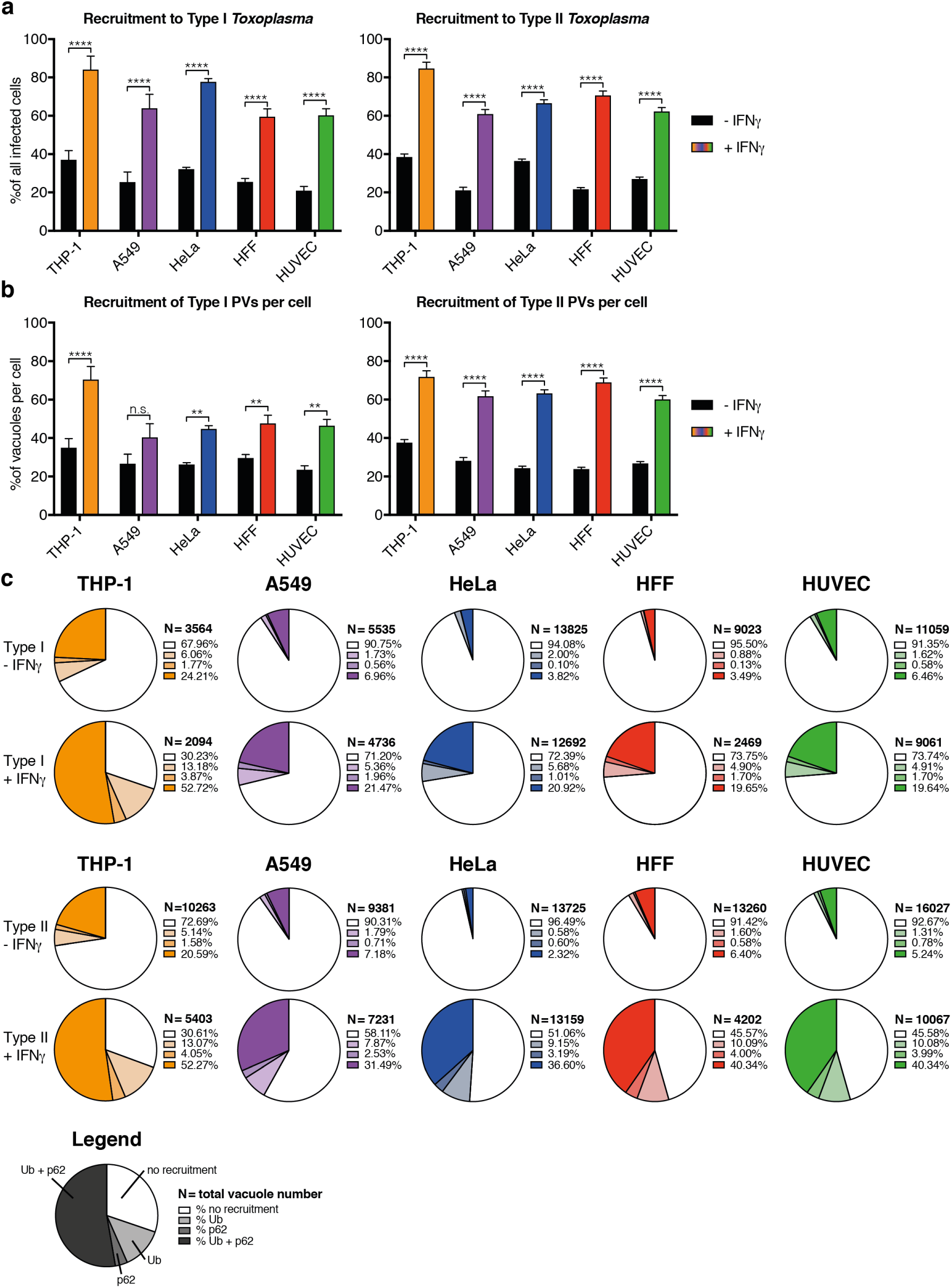
Ubiquitin and p62 host protein recruitment to *Toxoplasma gondii* type I and II vacuoles in 5 IFNγ-treated human cell lines at 6h post-infection. Cellular response to type I (RH) and type II (Pru) *Toxoplasma gondii (Tg)* infection. Percentage of infected cells that respond to *Tg* infection by decorating at least one vacuole with either ubiquitin, p62 or both. Proportion of vacuoles one cell can decorate with ubiquitin or p62 or both simultaneously. (c) Depicted are the average percentages *Tg* vacuoles decorated with host protein. Exact proportions can be found in the legend. The number of vacuoles analyzed is indicated. All data shown above represents the mean of N = 3 experiments ±SEM. Significance was determined using unpaired t-tests, n.s. = not significant, * p *≤* 0.0332, ** p *≤* 0.0021, *** p *≤* 0.0002 and **** p *<* 0.0001.

**Fig. S8:**
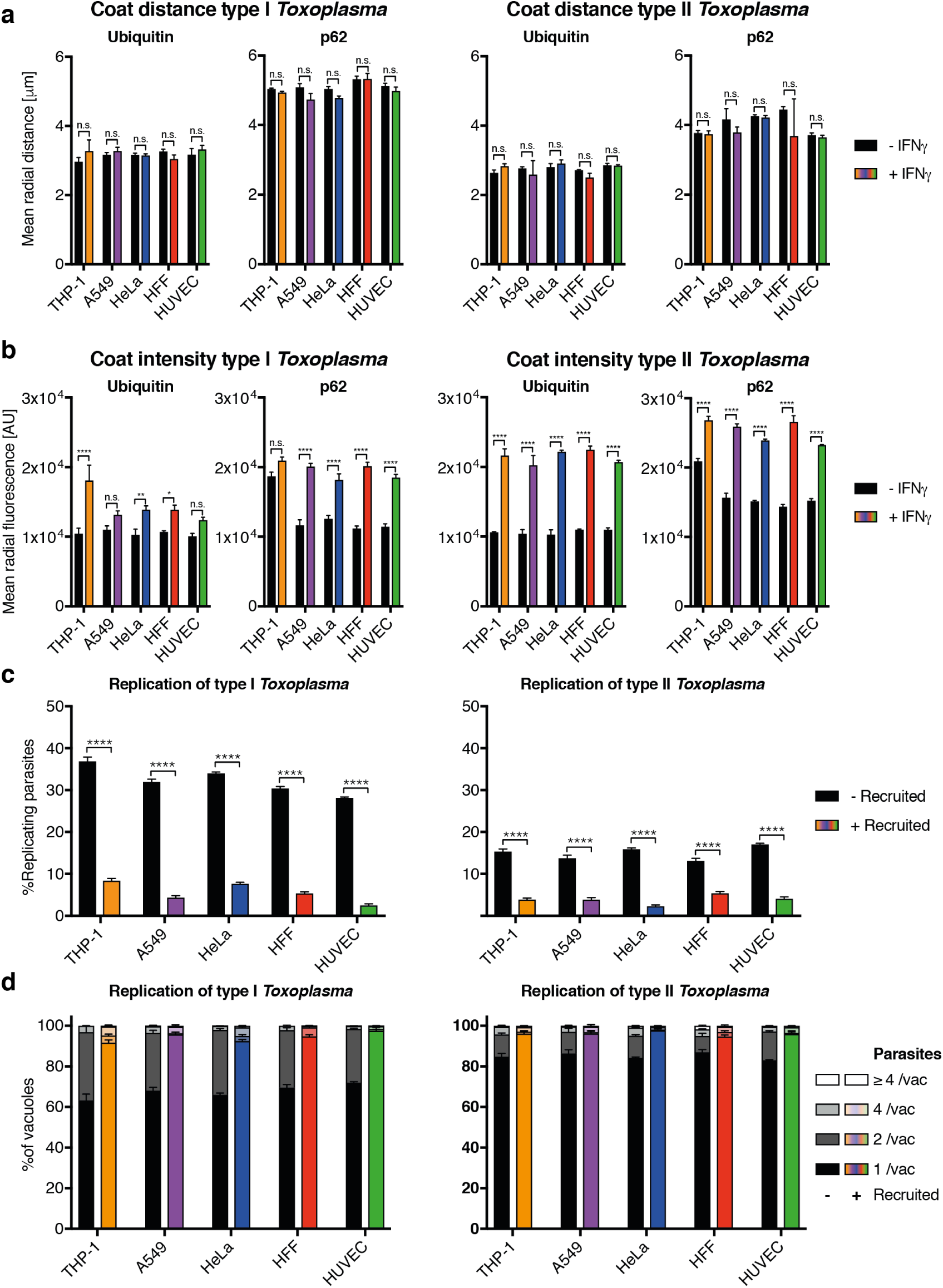
Characterization of the effect of host protein coating of *Toxoplasma gondii* type I and II vacuoles in 5 IFNγ-treated human cell lines at 6h post-infection. (a-b) Radial fluorescence intensity of host proteins around decorated type I (RH) and type II (Pru) *Toxoplasma gondii (Tg)* vacuoles. (a) Analysis of the coat distance to the centroid of *Tg* for ubiquitin and p62. (b) Intensity of the ubiquitin and p62 stain at the *Tg* vacuole. Significance was determined using unpaired t-tests, n.s. = not significant, * p ≤0.0332, ** p ≤0.0021, *** p ≤0.0002 and **** p *<* 0.0001. (c-d) Fate of *Tg* vacuoles grouped based on host protein recruitment. (c) Combined replication and recruitment analysis for non-decorated *Tg* vacuoles versus vacuoles co-decorated with ubiquitin and p62. (d) Replication distribution of *Tg* parasites contained in vacuoles with or without ubiquitin and p62 decoration. Significance was determined using 2-way ANOVA, n.s. = not significant, * p *≤* 0.0332, ** p *≤* 0.0021, *** p *≤* 0.0002 and **** p *<* 0.0001. All data shown above represents the means of N = 3 experiments ±SEM.

### Supplementary Table

**Table S1.** Overview and evaluation of existing software packages for analysis of fluorescence images in HCI experiments.

| Program | Reference | Machine learning and deep learning features and user friendliness |
| --- | --- | --- |
| Cellprofiler Analyst (CA) | Carpenter et al., 2006 | Deep learning (DL) is not a native part of CA. Using it requires writing python code. Furthermore, due to necessity to maintain the integration between DL framework (e.g. Google TensorFlow) and CA such code is even more complex defeating the purpose of the framework for DL application. |
| WND-CHARM | Orlov et al., 2008 | This collection of scripts requires coding and integration with up-to date libraries. |
| CellClassifier | Rämö, Sacher, Snijder, Begemann, & Pelkmans, 2009 | Being a milestone research contribution, CellClassifier is a discontinued legacy solution, making it currently unusable. |
| CellCognition (CC) | Held et al., 2010 | CC is a SVM-based user-friendly machine learning classification tool. However, similar to CA, DL is not a native part of CC, requires coding and comes with compatibility overhead. |
| Enhanced Cell Classifier (ECC) | Misselwitz et al., 2010 | While employing machine Learning (ML), ECC uses simplistic Support Vector Machine (SVM) classifier, which is not on par with modern machine learning techniques. ECC is a legacy solution. |
| Ilastik | Sommer, Straehle, Kothe, & Hamprecht, 2011 | Ilastik is ML-first oriented image analysis software with a great training interface and graphic processing unit (GPU) acceleration. It provides use of Random Forest ML classifier and SVM but has no DL support. |
| PhenoRipper | Rajaram, Pavie, Wu, & Altschuler, 2012 | PhenoRipper provides a lightweight analytical alternative to conventional computer vision approaches. However, it does not include the latest methodology like DL. |
| cellXpress (CX) | Laksameethanasan, Tan, Toh, & Loo, 2013 | CX offers great performance based on conventional computer vision algorithms. It is equipped with principle component analysis and SVM-based machine learning, however has no DL integration available. |
| Cytomine | Marée & et al., 2013 | Cytomine is a great implementation of python basic ML libraries. However, it is based on the Scikit-learn library, which is missing DL. Furthermore, it is cloud only, which is impractical for large-scale HCS datasets and/ or unpublished data. |
| PhenoDissim | Zhang & Boutros, 2013 | PhenoDissim is a tool to analyze phenotypic dissimilarity based on SVM, this approach has not been followed up with more advance ML techniques like DL. |
| BioConductor (BC) | Huber et al., 2015 | BC is an important tool in Bioinformatics with a substantial bioimage informatics capabilities. However, using BC requires coding. Furthermore, being aimed for R rather than python it provides poor integration with modern day DL libraries. |
| CP-CHARM | Uhlmann, Singh, & Carpenter, 2016 | Elaborated pure code analysis solution. However, requires understanding the code and integration with modern day libraries. |
| Advanced Cell Classifier (ACC) | Piccinini et al., 2017 | ML in ACC solution is based on a simplistic multi-layer perceptron network (MLP), which has been shown to be incapable of high complexity learning. |
| HTX | Arteta et al., 2017 | Great tool combining DL and graphical user interface in Matlab, however it lacks specific host-pathogen analysis. Furthermore, Matlab usage requires additional commercial licenses for highly specific tool-boxes used here, which significantly limits applicability of this framework. |
| Trainable Weka Segmentation | Arganda-Carreras et al., 2017 | Weka has arguably the largest ML classifiers collection, however for image analysis it relies on either ImageJ or KNIME integration and in absence of GPU acceleration no DL integration. |

